# Improved prediction of fungal effector proteins from secretomes with EffectorP 2.0

**DOI:** 10.1101/250464

**Authors:** Jana Sperschneider, Peter N. Dodds, Donald M. Gardiner, Karam B. Singh, Jennifer M. Taylor

**Author notes:** Author for correspondence: Jana Sperschneider.

## Abstract

Plant-pathogenic fungi secrete effector proteins to facilitate infection. We describe extensive improvements to EffectorP, the first machine learning classifier for fungal effector prediction. EffectorP 2.0 is now trained on a larger set of effectors and utilizes a different approach based on an ensemble of classifiers trained on different subsets of negative data, offering different views on classification. EffectorP 2.0 achieves accuracy of 89%, compared to 82% for EffectorP 1.0 and 59.8% for a small size classifier. Important features for effector prediction appear to be protein size, protein net charge as well as the amino acids serine and cysteine. EffectorP 2.0 decreases the number of predicted effectors in secretomes of fungal plant symbionts and saprophytes by 40% when compared to EffectorP 1.0. However, EffectorP 1.0 retains value and combining EffectorP 1.0 and 2.0 results in a stringent classifier with low false positive rate of 9%. EffectorP 2.0 predicts significant enrichments of effectors in 12 out of 13 sets of infection-induced proteins from diverse fungal pathogens, whereas a small cysteine-rich classifier detects enrichment only in 7 out of 13. EffectorP 2.0 will fast-track prioritization of high-confidence effector candidates for functional validation and aid in improving our understanding of effector biology. EffectorP 2.0 is available at http://effectorp.csiro.au.

## Introduction

Fungal pathogens have been estimated to cause annual crop yield losses of 15-20% and are a major threat to food security (Fischer *et al.,* 2014; Figueroa *et al.,* 2017). Fungi colonize plants through diverse infection structures and the use of toxic fungal secondary metabolites and secreted effector proteins that alter host cell structure and function, suppress plant defence responses or modulate plant cell physiology (Kamoun, 2006; Lo Presti *et al.,* 2015). Effectors are used by plant-pathogenic fungi to promote virulence and by symbiotic fungi to allow them to colonize their hosts. Fungal effectors can be attached to the fungal cell wall, can function in the plant apoplast or can translocate into plant cells where they might target specific host proteins or enter subcellular compartments (Lo Presti *et al.,* 2015). Accurate effector mining from genomic sequences is crucial to subsequent experimental validation and effector identification can enable disease control strategies. For example, effectors can be directly used in effector-assisted breeding to select plant lines with distinct recognition traits (Vleeshouwers & Oliver, 2014) and identification of both effectors and their targets could allow ‘decoy engineering’, where effector targets are fused as baits to a plant immune receptor to make an integrated ‘effector trap’ (Ellis, 2016).

Recent progress in big data genomics has resulted in many high-quality fungal pathogen genomes and gene expression profiles during plant infection, but accurate effector prediction methods are needed to harness the potential of these resources. The set of secreted proteins expressed during infection is typically too large for experimental investigation and contains many secreted non-effectors that play roles in niche colonisation and protection from competing microbes, differentiation of fungal structures and cell-to-cell communication (Rovenich *et al.,* 2014). Fungal effectors are diverse in sequence and share no conserved sequence motifs or obvious commonalities apart from their secretion from pathogen to the host. This lack of apparent unifying sequence-based features has led to ad-hoc fungal effector prediction approaches that are based on various combinations of characteristics observed in known effectors, such as a small protein size, a high cysteine content, evidence of diversifying selection, the genomic location of the gene in fast-evolving regions or gene expression *in planta* (Sperschneider *et al.,* 2015a). Including only a few features in effector prediction, such as the requirement of a small protein size, typically results in many false positive predictions and often overwhelmingly large effector candidate sets, such as 1,088 to 2,092 effector candidates predicted in stripe rust (Petre *et al.,* 2014). However, including additional features associated with effectors will capture only a small subset as none of these signals are common to all effectors. For example, some fungal effectors are highly enriched in cysteines whereas others do not feature any cysteines in their sequence and fungal effectors also vary in size. For example, the *Pyrenophora tritici-repentis* ToxB effector has 87 amino acids (aas) with four cysteines and is thought to function in the plant apoplast (Figueroa *et al.,* 2015), whereas the *Melampsora lini* AvrM effector has a sequence length of 314 aas and only one cysteine and acts intracellularly (Catanzariti *et al.,* 2006). However, a high cysteine content or small protein size alone also does not allow for accurate discrimination of apoplastic effectors from cytoplasmic effectors in fungi (Sperschneider, J. *et al.,* 2017). Taken together, the use of predefined criteria for effector prediction inherits the individual researcher’s potentially biased view of effector characteristics and is unable to uncover novel effectors with diverse characteristics.

An alternative approach is to use data to learn which features are important for effector prediction, rather than setting predefined criteria. This is achieved with machine learning, a family of statistical learning methods with the ability to identify patterns in data and recognise a particular class based on its features in observed data. Models trained on data sets of positive and negative classes are then applied to identify new instances of the class in unseen data. This data-driven approach has the capacity to identify new features not apparent to manual inspection and provide probabilistic predictions based on combinations of features, which represent advantages over using predefined criteria with binary cut-offs. We recently pioneered such a machine learning approach for fungal effector prediction called EffectorP (Sperschneider *et al.,* 2016) and demonstrated that machine learning can accurately predict novel effectors with diverse characteristics from secretomes as well as their localization in the plant cell (Sperschneider, Jana *et al.,* 2017; Sperschneider, J. *et al.,* 2017). We showed that EffectorP 1.0 was able to learn ‘effector rules’ from positive and negative training examples without having to apply user-chosen thresholds (Sperschneider *et al.,* 2016). EffectorP relies on fungal effectors as the positive training set and secreted non-effectors as the negative set. One limiting factor is that the negative training set consists of both undiscovered effectors and secreted non-effectors and therefore poses an unlabelled data classification problem. Furthermore, the positive training set used in EffectorP 1.0 was small and additional effectors are now available for inclusion in training. This has the potential to improve accuracy and will enable us to re-evaluate the ability of machine learning to accurately predict fungal effectors.

## Results

### Training of the ensemble classifier EffectorP 2.0

EffectorP 1.0 is a Naïve Bayes classifier that was trained on a positive training set of 58 experimentally supported fungal effectors from 16 fungal species. Since its development, additional fungal effectors have been described and for EffectorP 2.0 we used an expanded training set of 94 secreted fungal effectors from 23 species (Table 1). EffectorP 1.0 predicts 73% of the unseen effectors correctly, which demonstrate its ability to identify novel effectors, but also leaves room for improvement. We set out to investigate if re-training of EffectorP would improve prediction accuracy.

**Table 1:**
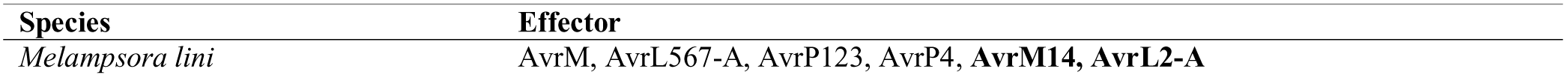
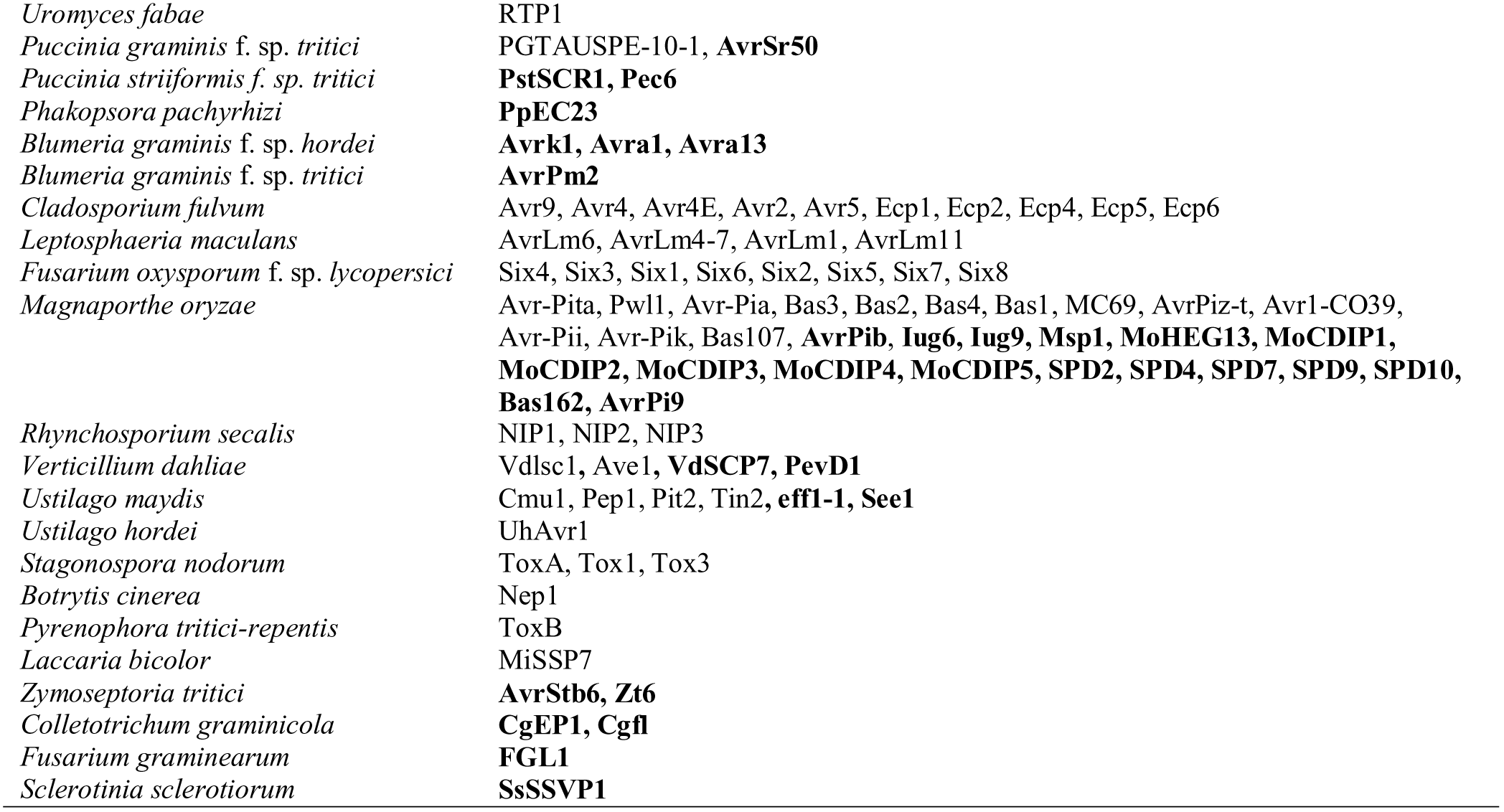
The set of fungal effector proteins used as positive training data. 94 fungal effectors were collected from the literature if they have experimental support and do not share sequence homology. Effectors that are not part of the EffectorP 1.0 training set are marked in bold. All sequences are available at http://effectorp.csiro.au/data.html.

EffectorP 1.0 was trained on a negative set consisting of predicted secreted proteins from the same pathogen species as the known effectors. Thus the negative training set includes both undiscovered effectors and non-effectors and therefore poses an unlabelled data classification problem. Whilst Naïve Bayes classifiers are fairly robust to unlabelled data classification and can tolerate noisy data (Bing *et al.,* 2003), other machine learning classifiers might not be able to learn effectively from such sets. To improve predictions, we collected three different subsets of negative training data that are less likely to contain positive instances, i.e. fungal effectors. First, secretomes were predicted from the same fungal pathogen/symbiont species that were used in the positive set if they have a publicly available predicted gene set (Table 1). The combined secretome was homology-reduced and this resulted in a filtered predicted pathogen secretome of 11,277 proteins. This set will contain both undiscovered effectors and secreted non-effectors, which poses a challenge for machine learning classifiers that traditionally learn from labelled data. Therefore, we applied EffectorP 1.0 to exclude predicted effectors from the secretomes (*n =* 6,138). We also collected homology-reduced sets of secreted fungal proteins from fungi not pathogenic on plants, namely from 27 saprophyte secretomes (*n =* 12,939) and from 10 animal-pathogenic fungal secretomes (*n =* 2,763). These sets are less likely to contain plant pathogenic effectors and were not filtered for EffectorP 1.0 predicted effectors.

As we have large amounts of negative training data *(n =* 21,840), we used an ensemble learning approach of classifiers that each take a different subset of negative training data and thus provide a different view on classification (Fig.1). Overall, we chose a total of 50 best-performing models comprised of: ten Naïve Bayes classifiers and ten C4.5 decision trees that discriminate between fungal effectors and secreted pathogen proteins; ten Naïve Bayes classifiers and ten C4.5 decision trees that discriminate between fungal effectors and secreted saprophyte proteins; and five Naïve Bayes classifiers and five C4.5 decision trees that discriminate between fungal effectors and secreted animal pathogen proteins. In 10-fold cross validation, the Naïve Bayes classifiers achieved on average high sensitivity, whereas the C4.5 decision trees have high specificity (Supporting Information Table S2). To generate EffectorP 2.0, we combined these 50 models into an ensemble classifier to utilize their distinct prediction strengths (Fig. 1). Each model has seen a different subset of negative training data and for a given protein sequence input returns a probability that it is an effector or a non-effector. EffectorP 2.0 returns a final prediction using a voting approach on the predicted probabilities of each model. A protein is classified as an effector if the average probability for the class ‘effector’ is higher than the average probability for the class ‘non-effector’.

**Fig. 1:**
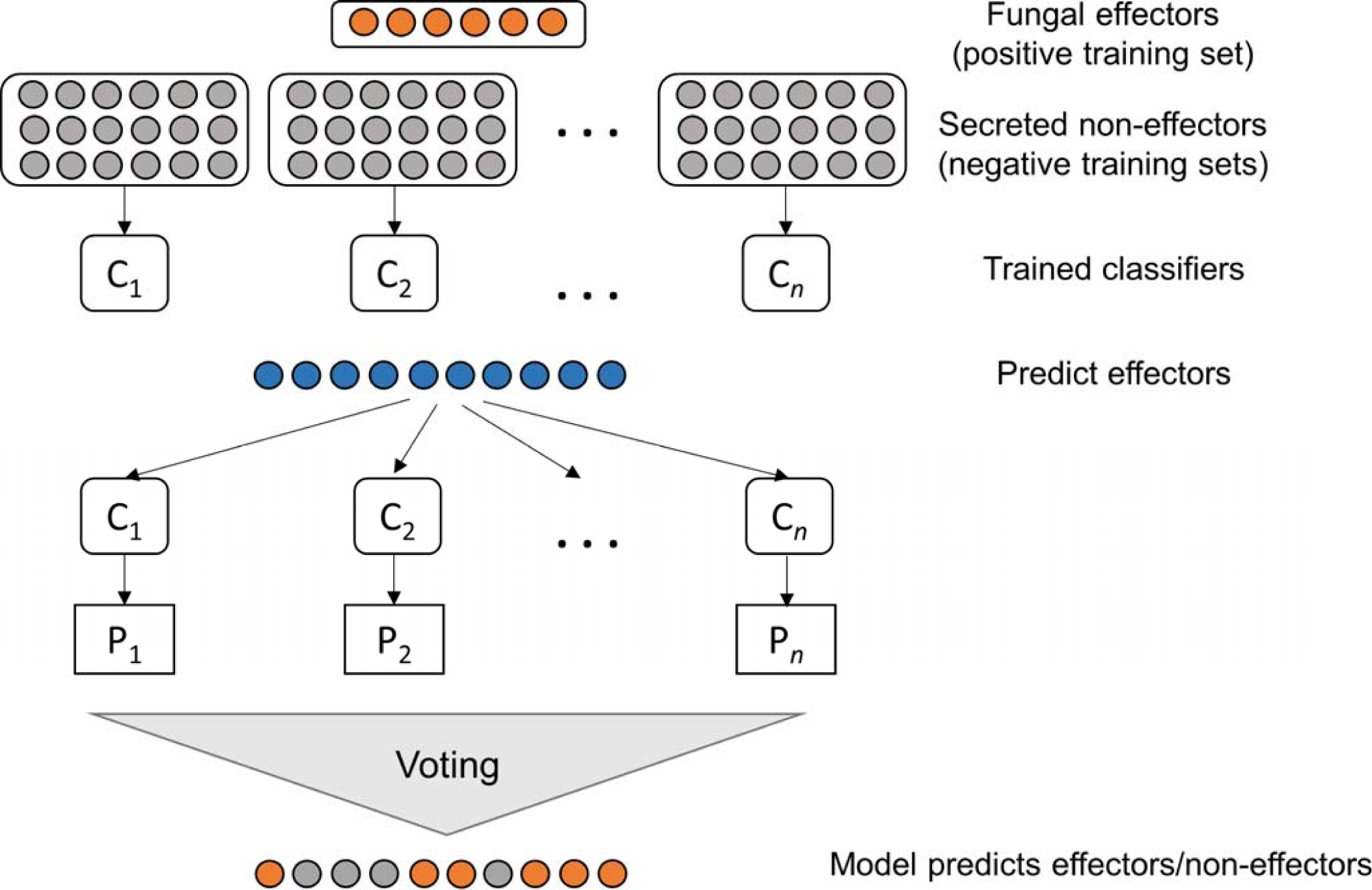
Workflow for the EffectorP 2.0 classifier that combines an ensemble of machine learning classifiers. Each classifier C_i_ has seen a different subset of the negative training data and predicts effectors in unseen data with probability P_i_. The probabilities are combined into an overall vote on whether an unseen protein is an effector or non-effector.

### Influential features for effector prediction include protein size, protein net charge as well as the amino acids serine and cysteine

To detect the most discriminative features in the EffectorP 2.0 classification, we analyzed the distribution of features for the proteins employed in training of all 50 models. Four features were found to be different at a significance threshold of *p <* 10^−5^ in distribution between the positive sequence set (effectors) and the negative sequence set (proteins labelled as non-effectors) (Fig. 2). Differences in feature distribution for these four features were also previously reported in the EffectorP 1.0 model as particularly striking (Sperschneider *et al.,* 2016), confirming their importance in fungal effector classification. As a group, the effectors exhibit lower molecular weight, a higher percentage of cysteines (C) and a lower percentage of serines (S) than the proteins in the negative sequence set. The distribution of protein net charge for effectors occupies a narrow range around neutral to slightly positive (Fig. 2). We also found significant differences *(p <* 0.05) in distribution between effectors and the negative sequence set for additional features (Fig. 2). These are depletion in aliphatic amino acids, leucine (L), proline (P), threonine (T), tryptophan (W), disorder propensity and bulkiness as well as enrichment in basic amino acids, interface propensity, glycine (G), lysine (K) and asparagine (N) for effectors. Only enrichment in tryptophan content in effectors was also reported in the EffectorP 1.0 model.

**Fig. 2:**
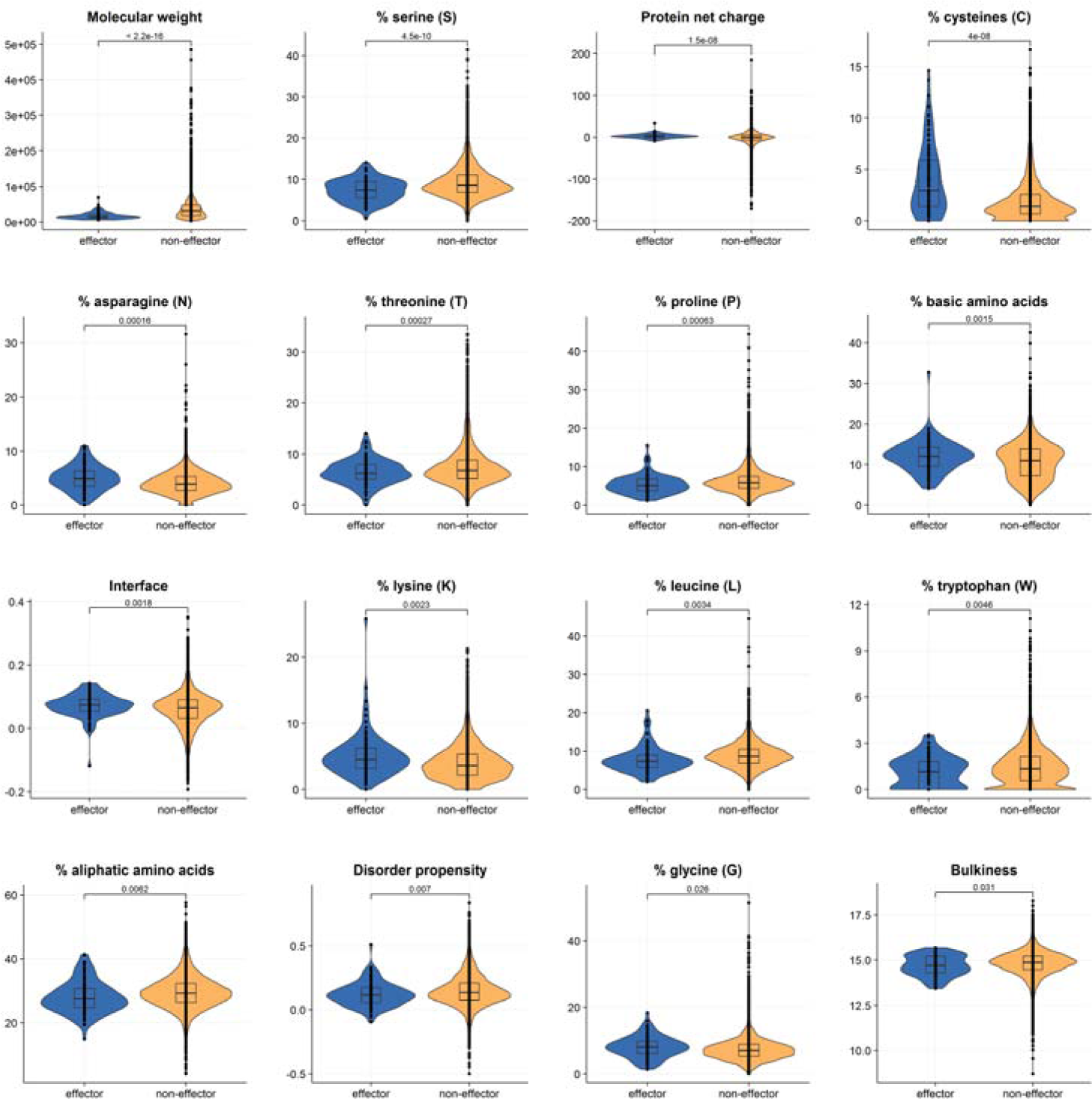
The most influential features in effector prediction appear to be a small protein size, low serine content, a protein net charge around the neutral range and a high cysteine content. Significant differences *(p <* 0.05) in distribution between effectors and the negative sequence set for additional features were also observed. These are depletion in aliphatic amino acids, leucine (L), proline (P), threonine (T), tryptophan (W), disorder propensity and bulkiness as well as enrichment in basic amino acids, interface propensity, glycine (G), lysine (K) and asparagine (N) for effectors. Extreme outliers in the protein net charge plot were removed for clarity (full figure given in Supporting Information File S1). All data points were drawn on top of the box plots as black dots. Significance between groups is shown as horizontal brackets and was assessed using *t*-tests. The lower and upper hinges correspond to the first and third quartiles and the upper (lower) whiskers extend from the hinge to the largest (smallest) value that is within 1.5 times the interquartile range of the hinge. Data beyond the end of the whiskers are outliers.

Machine learning can be a black box learning process where the reasons for an individual prediction are hidden. However, C4.5 decision trees are white box models and their decision making process is transparent through navigation along tree branches. As examples, we plotted two of the 10 C4.5 decision trees that discriminate between fungal effectors and secreted pathogen proteins (Supporting Information Figs. S1 and S2). This demonstrates that the decision tree classifiers use a complex set of features and not only the most discriminative features (protein size, protein net charge as well as the amino acids serine and cysteine) for effector classification. In particular, the decision tree in Figure S2 does not utilize serine content as a feature in classification and still achieves high classification accuracy. Taken together, this analysis confirms the importance of specific combinations of features as previously found in the EffectorP 1.0 model, but also illustrates that accurate fungal effector prediction machine learning classifiers rely on a diverse set of features.

### EffectorP 2.0 improves fungal effector prediction accuracy from secretomes

Machine learning classifiers can overfit/overtrain to memorize the training data which leads to low accuracy on unseen data. Therefore, independent test sets are important to estimate prediction ability. We collected independent positive and negative test sets to assess the performance of EffectorP 2.0. To estimate the false positive rate, we first used fungal, plant and mammalian proteins with a predicted signal peptide that are not extracellular (localization to ER, Golgi or membranes or with GPI-anchors). A low false positive rate on these proteins ensures that EffectorP is not merely predicting the presence of a signal peptide. We also used secreted saprophyte proteins as well as fungal proteins from PHI-base (Urban *et al.,* 2017) that are annotated as having an unaffected pathogenicity phenotype. Whilst proteins with an unaffected pathogenicity phenotype are not necessarily non-effectors, we expect to see a low percentage of predicted effectors. A simple classifier based on a small protein size (<= 300 aas) has a false positive rate of 40.4% on these three sets. A small, cysteine-rich classifier (<= 300 aas, >= 4 cysteines) has a false positive rate of 19% and EffectorP 1.0 has a false positive rate of 18.3%. EffectorP 2.0 has the lowest false positive rate of 11.2% (Table 2). A combination of EffectorP 1.0 and 2.0, where a protein is a predicted effector only if both classifiers label it as an effector achieves the lowest false positive rate of 9.4%.

**Table 2:** Independent validation of EffectorP’s prediction accuracy.

| Data set | # of proteins | Predicted effectors |  |  |  |  |
| --- | --- | --- | --- | --- | --- | --- |
|  |  | EffectorP 2.0 | EffectorP 1.0 | EffectorP 1.0 and 2.0 | Small size classifier | Small, cysteine-rich classifier |
| Fungal saprophyte secreted proteins | 24,432 | 2,865 (11.7%) | 4,774 (19.5%) | 2,444 (10%) | 10,529 (43.1%) | 4,961 (20.3%) |
| Fungal, plant and mammalian proteins with signal peptide and localization to ER, Golgi, membranes or with GPI-anchors | 2,631 | 220 (8.4%) | 294 (11.2%) | 164 (6.2%) | 654 (24.9%) | 307 (11.7%) |
| Fungal proteins with unaffected pathogenicity phenotype | 938 | 45 (4.8%) | 59 (6.3%) | 36 (3.8%) | 128 (13.6%) | 60 (6.4%) |
|  | **28,001** | **3,130 (11.2%)** | **5,127 (18.3%)** | **2,644 (9.4%)** | **11,311 (40.4%)** | **5,328 (19%)** |
| Fungal effector positive training set | 94 | 89 (94.7%) | 80 (85.1%) | 79 (84%) | 88 (93.6%) | 53 (56.4%) |
| Fungal effector independent test set | 21 | 16 (76.2%) | 16 (76.2%) | 16 (76.2%) | 19 (90.5%) | 10 (47.6%) |
| **Accuracy** |  | **88.8%** | **81.7%** | **90.5%** | **59.8%** | **80.9%** |

To assess false negative predictions, we also applied these predictors to the training data of 94 fungal effectors (Table 2). EffectorP 2.0 only predicts five of these proteins as non-effectors: the *Phakopsora pachyrhizi* effector PpEC23, the *Blumeria graminis* f. sp. *hordei* effector Avrk1, the *Magnaporthe oryzae* effector MoCDIP2, the *Ustilago maydis* effector eff1-1 and the *Colletotrichum graminicola* metalloproteinase effector Cgfl. This is an improvement on Effector P1.0, which correctly predicted only 80 of the 94 positive examples. However, it is also important to assess overfitting on training data and use unseen fungal effectors independent from the training set for validation of the estimated true positive rate. Therefore we collected 21 effectors (Table 3) that either share sequence similarity with an effector in the training set and were therefore eliminated in the homology reduction step (Mg3LysM, BEC1054, BEC1011, AvrLm2) or were overlooked during initial literature searches for training the EffectorP 2.0 model (SAD1, CSEP-07, CSEP-09, SIS1, CSEP0055, BEC1019, Bcg1, CSEP0105, CSEP0162, AvrLmJ1, AvrLm3, XylA, Ecp7, PIIN08944, FGB1, AvrPm3, AvrSr35). On this independent test set, both EffectorP 1.0 and 2.0 show equal performance and correctly predict 76.2% of effectors (Table 2, 3). On the total set of 115 effectors, the small size classifier correctly predicts 93% of effectors, but the small, cysteine rich classifier only correctly predicts 54.8% of effectors. On the combined positive and negative sets, EffectorP 2.0 has the highest accuracy of 88.8% out of the four single classifiers. The simple classifier based on a small size has the lowest accuracy of 59.8%, largely due to its high false positive rate (Table 2). The combined EffectorP 1.0/2.0 classifier achieves the highest accuracy of 90.5% due to its low false positive rate. Whilst the combined EffectorP 1.0/2.0 classifier misses more effectors than EffectorP 2.0 or 1.0, it is a highly stringent method for predicting effectors in secretomes. In the following, we assess the prediction abilities of EffectorP 1.0 compared to 2.0 in more detail.

**Table 3:**
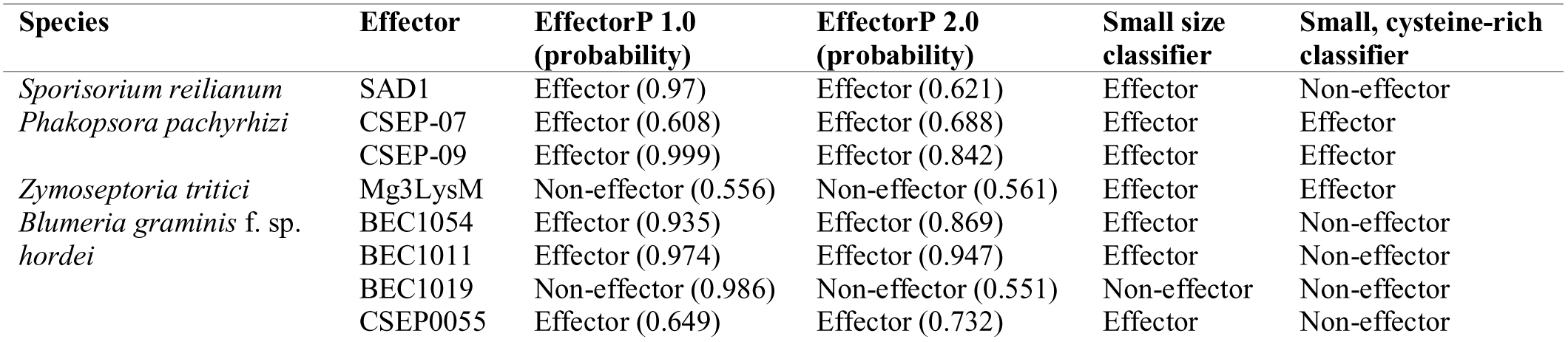
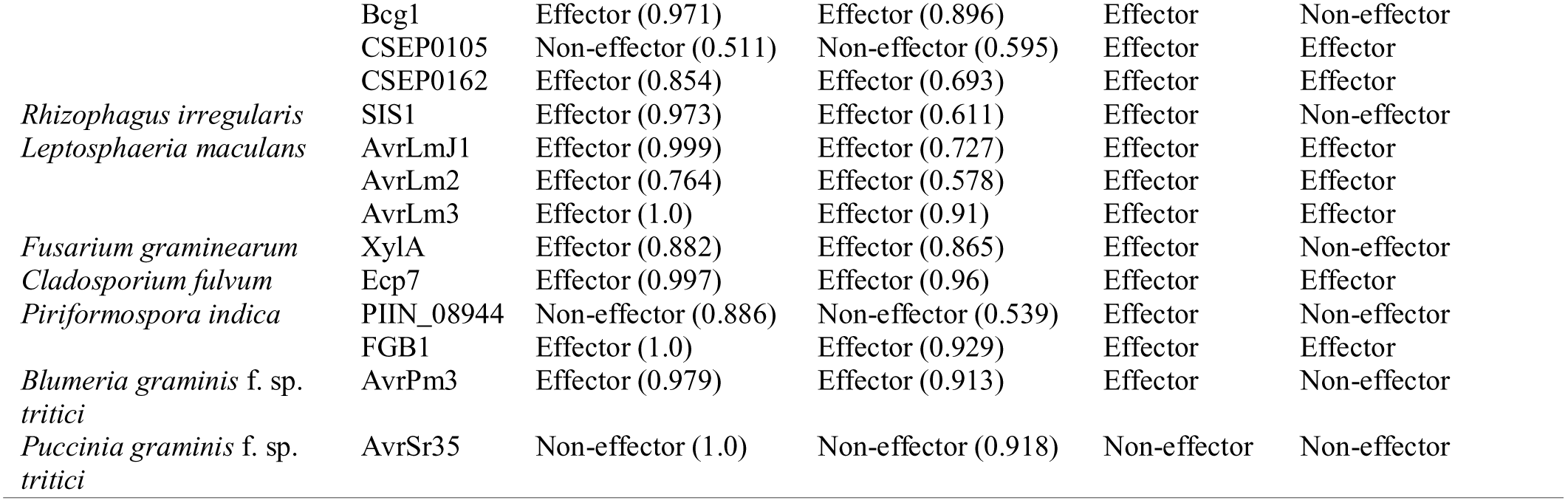
Independent test set of fungal effectors that were not used in training of EffectorP 2.0.

### Sets of infection-induced proteins are enriched for effectors predicted by EffectorP 2.0

Effectors are often induced during infection and thus the set of genes differentially expressed during infection should be enriched for effectors. However, not all genes that are differentially expressed during infection encode effector proteins and therefore, sets of differentially expressed genes need to be filtered further to detect effectors. We collected 13 gene sets from the literature that were labelled as containing effector candidates based on their induction during infection as well as other criteria (Table 4). For example, a study by Germain *et al.* (2016) identified 16 candidate effectors from 1,184 small, secreted *Melampsora larici-populina* proteins. These 16 candidates were selected based on their expression in a haustoria-specific cDNA library and the transcriptome of laser microdissected, rust-infected poplar leaves as well as their small size of less than 300 aas. As another example, Kettles *et al.* (2017) selected 63 *Zymoseptoria tritici* candidate effectors on the basis of being induced during early wheat leaf infection leading up to the transition to the necrotrophic growth phase. In total, 4 of the 13 sets contain infection-induced effector candidates that were pre-selected based on a small size (<= 300 aas).

**Table 4:**
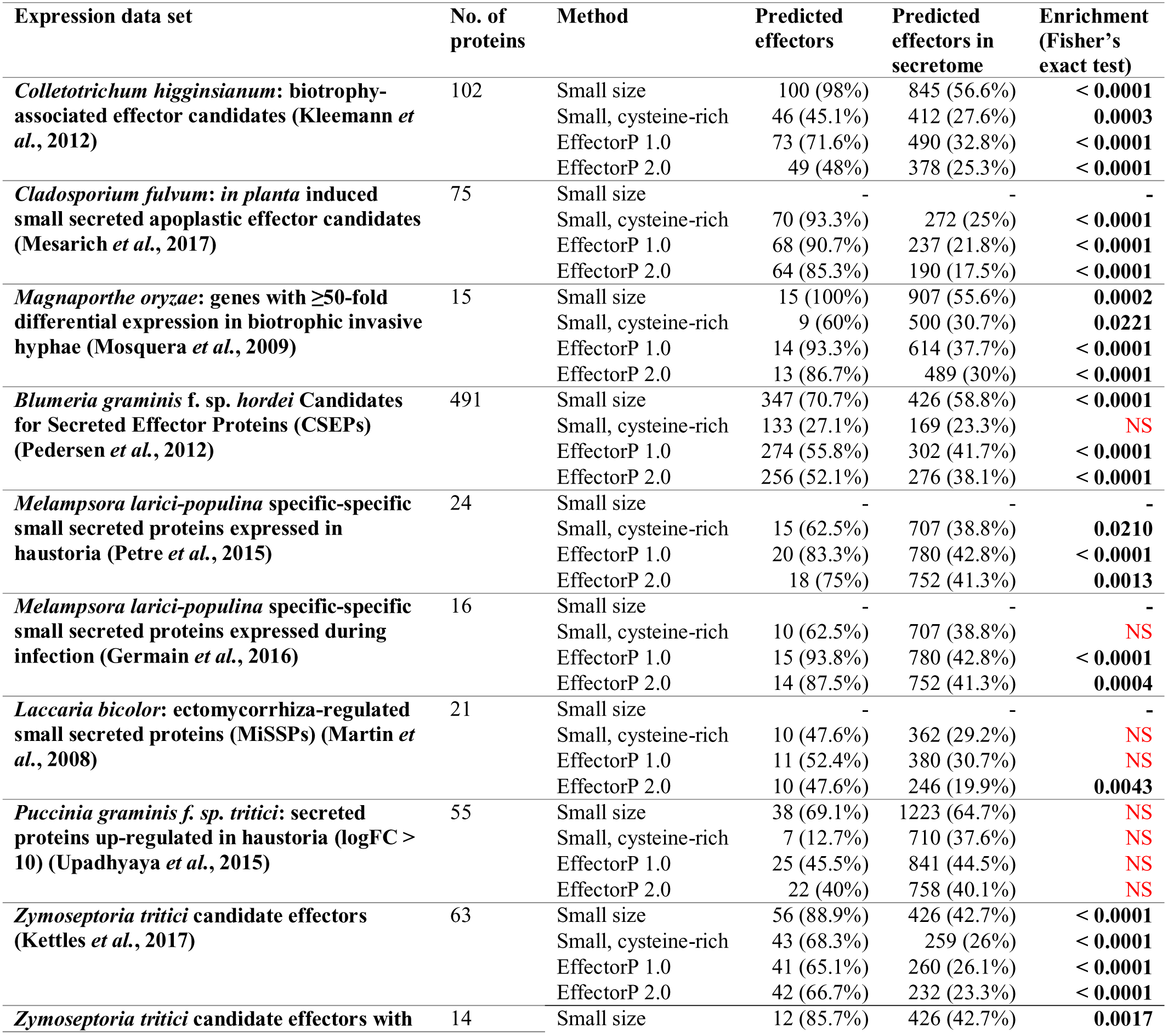
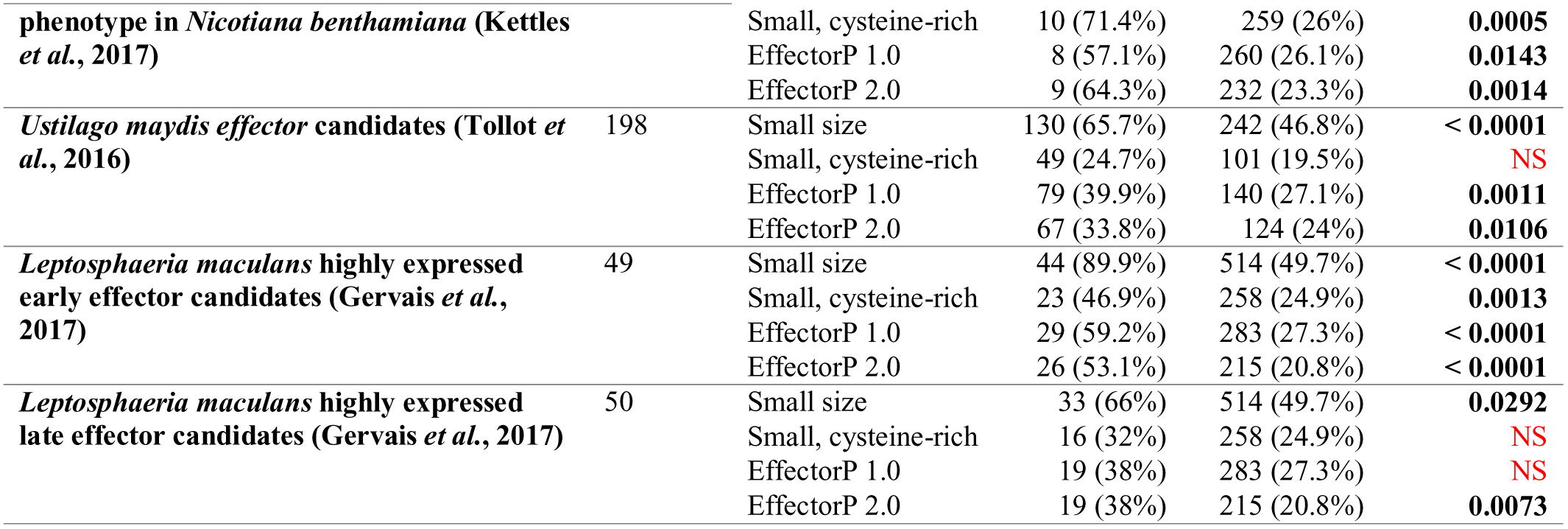
Enrichment of predicted effector candidates in expression data sets of early infection stages. For each expression data set, the percentage of predicted effector candidates by EffectorP is shown and compared to the percentage of predicted effector candidates in the secretome. The small size classifier is only applied to sets that are not pre-selected based on a small size.

We assessed whether the 13 sets containing infection-induced effector candidates are also enriched for effector candidates predicted by EffectorP 1.0 or 2.0, by a small size classifier or by a small, cysteine-rich classifier when compared to the whole secretome of each species. We did not test the small size classifier on sets containing effector candidates that were pre-selected based on a small size (<= 300 aas). We found significant enrichments for predicted effector candidates in 12 out of 13 sets (92.3%) using EffectorP 2.0 (Table 4). A small, cysteine-rich classifier only returns significant enrichments for predicted effectors in 7 out of 13 sets (53.9%) and EffectorP 1.0 in 10 out of 13 sets (76.9%). A small size classifier returns significant enrichments for predicted effectors in 8 out of 9 sets (88.9%). Surprisingly, we did not observe enrichment for predicted effectors with any of the four classifiers in secreted proteins of *P. graminis* f. sp. *tritici* highly up-regulated in haustoria compared to germinated spores (Table 4). This could indicate that rusts might utilize undiscovered effector proteins with different properties to the training set, such as effectors of larger size. This is supported by the recent discovery of AvrSr35, a 578 aas *P. graminis* f. sp. *tritici* effector protein (Salcedo *et al.,* 2017). Alternatively, haustorial secretomes might contain many non-effectors such as proteins involved in signalling or in incorporating nutrients from the host (Garnica *et al.,* 2014). Taken together, whilst effector function has not been established for all genes in these candidate sets, the enrichment for predicted effectors in infection-induced sets underlines the ability of EffectorP 2.0 to accurately predict unseen effectors.

### EffectorP 2.0 reduces the average number of effectors predicted for fungal plant symbionts and saprophytes by 40%

We tested EffectorP 2.0 on predicted secretomes from 93 fungal species, including pathogens and non-pathogens (Supporting Information Table S3) and recorded the percentages of secreted proteins that are predicted effectors (Supporting Information File S2). The highest proportions of predicted effectors were found in the obligate biotrophs *Melampsora laricis-populina* (41.3%), *Puccinia graminis* f. sp. *tritici* (40.3%), *Blumeria graminis* f. sp. *hordei* (38.1%) and *Puccinia striiformis f.* sp. *tritici* (37.6%). Amongst the fungal plant pathogens, the lowest proportions of predicted effectors were recorded for the necrotrophs *Heterobasidion annosum* (10.4%), *Sclerotinia sclerotiorum* (13.6%), *Botrytis cinerea* (13.7%) and *Penicillium digitatum* (13.9%). Necrotrophic pathogens utilize many secreted non-effectors, such as cell wall degrading enzymes, to kill and degrade the plant cell wall whereas biotrophic pathogens must suppress plant defence mechanisms through effectors.

On average, EffectorP 2.0 predicts that plant pathogen secretomes consist of 24.9% effectors and that saprophyte secretomes consist of 11.7% effectors (Tables 4 and 5). EffectorP 2.0 reduces the average number of predicted effectors in fungal plant symbiont and fungal saprophyte secretomes by over 40% when compared to EffectorP 1.0 (Table 5, Fig. 3). Both EffectorP 2.0 and EffectorP 1.0 also predict lower proportions of effectors for fungal animal pathogens than for fungal plant pathogens (Table 5), suggesting that effector repertoires of fungal animal pathogens are different to those of their plant-pathogenic counterparts. One notable exception is the secretome of *Enterocytozoon bieneusi,* an obligate intracellular parasite (49 predicted effectors, 36% of secretome predicted as effectors). Shortened protein-coding sequences due to genome compaction have been reported in *E. bieneusi* (Akiyoshi *et al.,* 2009) and might lead to higher than expected false positive predictions. Therefore, we also assessed effector prediction rates for small secreted proteins (< 300 aas) only. For plant pathogens, EffectorP 2.0 predicts that 47.8% of small secreted proteins are effectors, whereas for plant symbionts and saprophytes this is reduced to 29.9% and 26.3%, respectively. This underlines that EffectorP 2.0 is not selecting effectors based on a small size alone. Small secreted proteins in saprophytes are mostly functionally uncharacterized and might function in a variety of processes unrelated to plant-pathogen interactions. Compared to a small, cysteine-rich classifier EffectorP 2.0 predicts significantly lower proportions of effectors for plant symbionts and saprophytes, but not for plant pathogens (Fig. 3). This lack of correlation for all groups tested underlines that EffectorP 2.0 is not selecting effectors based on a small size and a high cysteine content alone and reflects the reduced false positive rate of EffectorP 2.0.

**Fig. 3:**
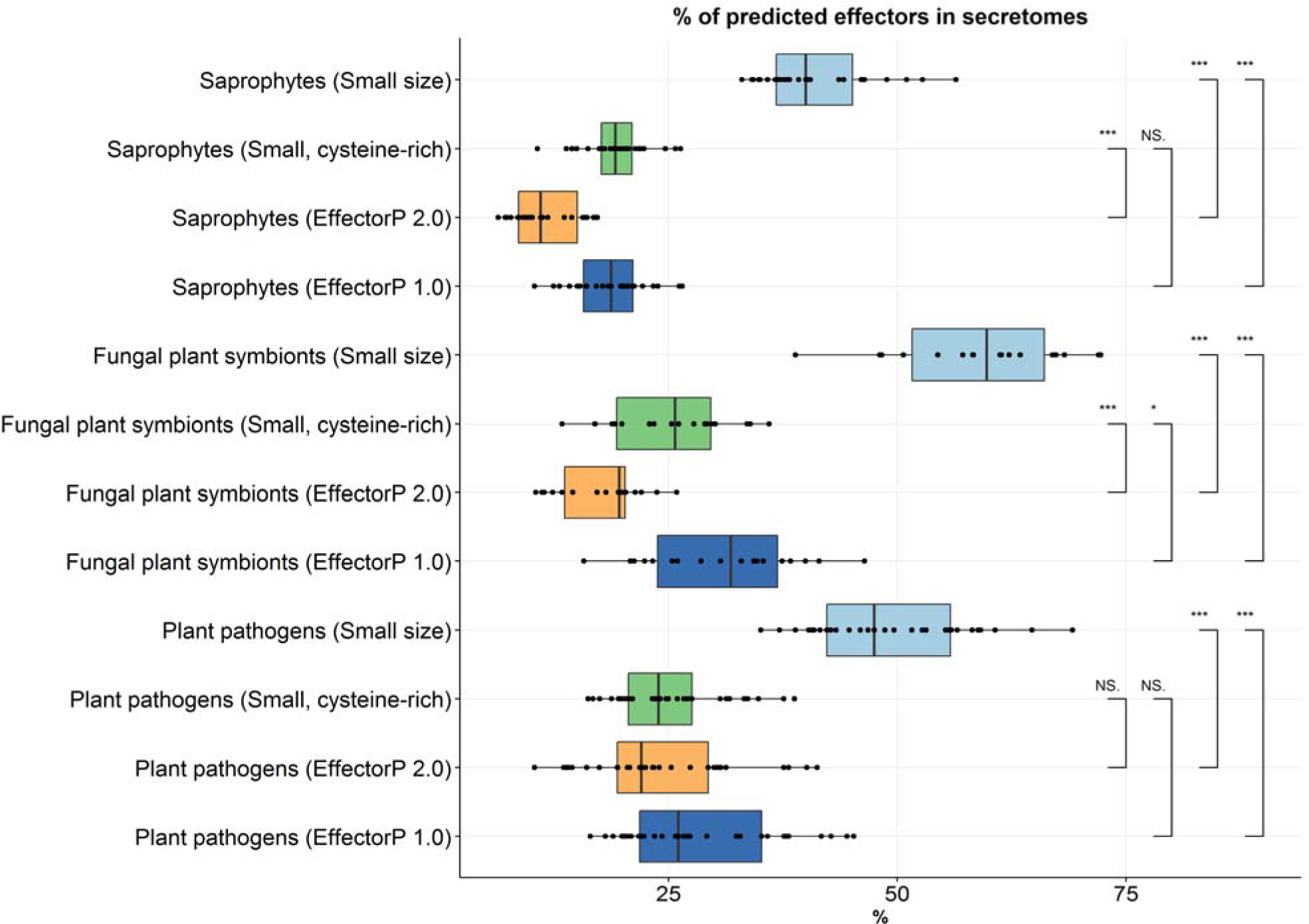
Proportions of predicted effectors in fungal secretomes using EffectorP 1.0, EffectorP 2.0, a small size classifier and a small, cysteine-rich classifier. All data points were drawn on top of the box plots as black dots. Significance between groups is shown as horizontal brackets and was assessed using f-tests (NS: not significant, * < 0.05, ** < 0.01 and *** < 0.001). The lower and upper hinges correspond to the first and third quartiles and the upper (lower) whiskers extend from the hinge to the largest (smallest) value that is within 1.5 times the interquartile range of the hinge. Data beyond the end of the whiskers are outliers.

**Table 5:** Predicted effectors in secretomes for groups of fungal species.

| Secretomes | Average of predicted effectors |  | % decrease in predicted effectors<br>(EffectorP 2.0 compared to EffectorP 1.0) |
| --- | --- | --- | --- |
|  | EffectorP 1.0 | EffectorP 2.0 |  |
| Plant pathogens | 338 (29.6%) | 284 (24.9%) | -16.0% |
| Fungal symbionts of plants | 305 (30.8%) | 177 (17.8%) | -42.0% |
| Fungal pathogens of animals | 108 (20.9%) | 83 (16.1%) | -23.2% |
| Saprophytes | 177 (19.5%) | 106 (11.7%) | -40.1% |

We then further investigated the properties of effectors that are only predicted by one of the versions of EffectorP, but not by the other, for all 93 secretomes (Table 6). Effector candidates predicted only by EffectorP 2.0 are on average of longer sequence length (n = 2,304, average sequence length 229 aas) than those that are only predicted by EffectorP 1.0 (n = 8,635, average sequence length 138 aas) or by both versions (n = 14,128, average sequence length 148 aas) (Fig. 4). Effector candidates predicted only by EffectorP 1.0 or 2.0 are lower in cysteine content compared to effector candidates predicted by both versions (Fig. 4). We then tested for enrichment and depletion of protein functional classes amongst the effector candidates predicted by EffectorP 1.0 and 2.0 from a total of 24,075 secreted proteins of the 21 plant pathogens (Table 1). The vast majority of effector candidates predicted by either EffectorP 1.0 or 2.0 are proteins without functional annotation. However, we observed that both sets of predicted effector candidates are enriched for proteins with pectate lyase activity, peptidyl-prolyl cis-trans isomerase activity and endopeptidase inhibitor activity (Table 6). Some proteins with peptidyl-prolyl cis-trans isomerase activity have been implicated to function as virulence factors (Unal & Steinert, 2014). EffectorP 2.0 predicted effectors are enriched for proteins involved in pathogenesis and defence response (Table 6). However, EffectorP 1.0 predicted effector candidates are also enriched for proteins that do not appear related to effector function or to secreted proteins, but rather to intracellular proteins (Table 6), and might reflect the higher false positive rate of EffectorP 1.0 as well as the false positive rate of the signal peptide prediction tools SignalP 3.0 and TargetP.

**Fig. 4:**
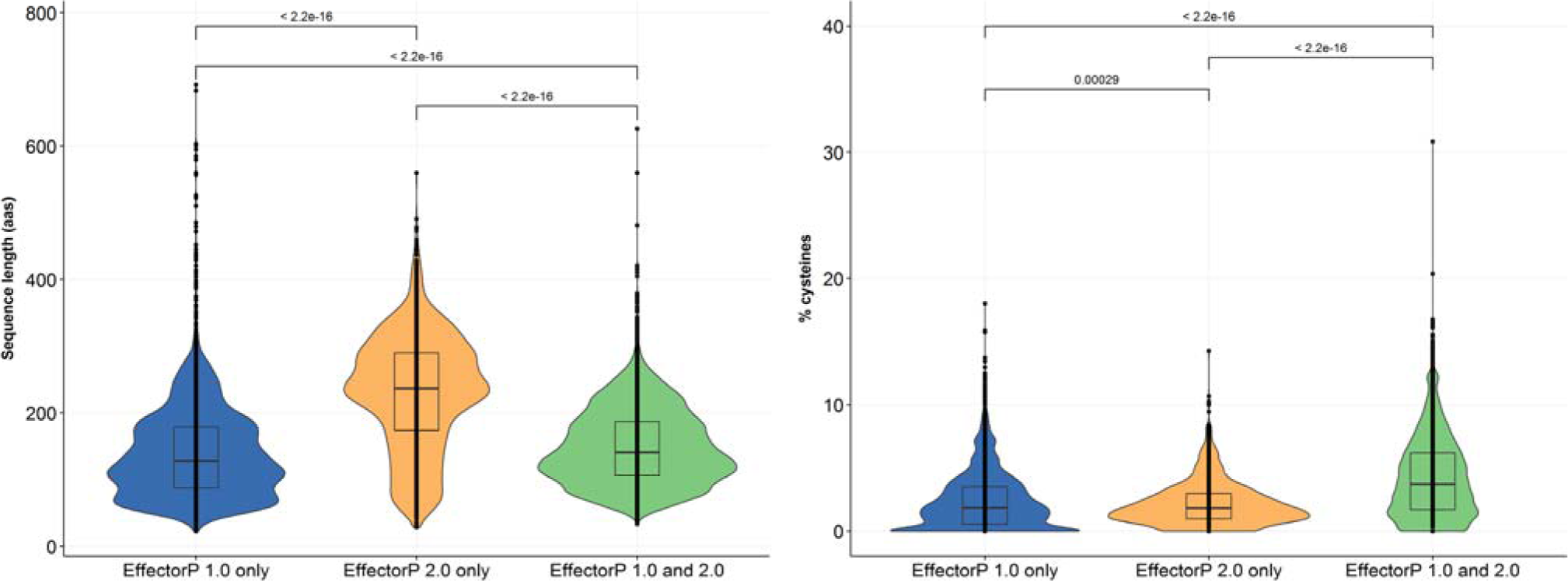
Differences in sequence length and cysteine content for effectors predicted by different versions of EffectorP. All data points were drawn on top of the box plots as black dots. Significance between groups is shown as horizontal brackets and was assessed using f-tests. The lower and upper hinges correspond to the first and third quartiles and the upper (lower) whiskers extend from the hinge to the largest (smallest) value that is within 1.5 times the interquartile range of the hinge. Data beyond the end of the whiskers are outliers.

**Table 6:** GO term enrichment analysis of predicted effector candidates.

| Comparison | Over-represented GO term description | # proteins in test set | # proteins in reference set | FDR |
| --- | --- | --- | --- | --- |
| **Test set:** EffectorP 2.0 predicted<br>**Reference set:** Secreted pathogen proteins | Pectate lyase activity | 26 | 11 | $2.9 \times 10^{-8}$ |
| | Peptidyl-prolyl cis-trans isomerase activity | 15 | 5 | $2.6 \times 10^{-5}$ |
|  | Pathogenesis | 12 | 10 | 0.02 |
|  | Defence response | 14 | 14 | 0.02 |
|  | Endopeptidase inhibitor activity | 7 | 3 | 0.03 |
| **Test set:** EffectorP 1.0 predicted<br>**Reference set:** Secreted pathogen proteins | Peptidyl-prolyl cis-trans isomerase activity | 18 | 2 | $1.9 \times 10^{-7}$ |
| | Inner mitochondrial membrane organization | 10 | 1 | $3.6 \times 10^{-4}$ |
| | Intracellular sterol transport | 8 | 0 | $5.7 \times 10^{-4}$ |
| | Fungal-type vacuole lumen | 8 | 0 | $5.7 \times 10^{-4}$ |
| | Pectate lyase activity | 22 | 15 | $8.4 \times 10^{-4}$ |
|  | Endopeptidase inhibitor activity | 9 | 1 | 0.001 |
|  | Chaperone-mediated protein folding | 7 | 0 | 0.001 |
|  | FK506 binding | 7 | 0 | 0.002 |
|  | Nuclear envelope organization | 6 | 0 | 0.002 |
|  | Regulation of COPII vesicle coating | 6 | 0 | 0.006 |
|  | Endoplasmic reticulum exit site | 6 | 0 | 0.006 |
|  | Mitochondrial inner membrane | 18 | 15 | 0.01 |
|  | Mitochondrial respiratory chain complex IV assembly | 5 | 0 | 0.02 |
|  | Mitochondrion morphogenesis | 5 | 0 | 0.02 |
|  | COPII vesicle coat | 6 | 1 | 0.03 |

## Discussion

Given the high diversity of fungal effectors, it seems an unexpected finding that a machine learning classifier can accurately distinguish diverse effectors from secreted non-effectors. However, classifiers such as decision trees can have multiple paths that lead to a prediction as an effector and one can speculate that different paths might relate to different classes of effectors, such as apoplastic or cytoplasmic ones. Decision trees can also learn feature interactions, whereas Naïve Bayes classifiers identify the importance of individual features, but not relationships amongst features. This might be advantageous for effector prediction, e.g. a Naïve Bayes classifier can learn that a small protein size or a high cysteine content is important for effectors, but it does not learn that proteins have to be small and at the same time cysteine-rich to be effectors. Unlike Naïve Bayes classifiers, decision trees are non-parametric, which gives them the ability to e.g. assign a very low protein size to non-effectors, a low to medium protein size to effectors and a large protein size to non-effectors. However, decision trees are prone to overfitting, especially on small training data sets, which can lead to a limited ability to correctly classify unseen data. Naïve Bayes classifiers can deliver robust performance on small training data sets and an ensemble classifier like EffectorP 2.0 is capable to draw on the strengths of both decision trees and Naïve Bayes classifiers.

On the current training set, low molecular weight is an important feature in fungal effector classification. However, it is possible that fungal pathogens employ classes of larger effector proteins that have thus far not been recognized. For example, the recently discovered *Puccinia graminis f.* sp. *tritici* effectors AvrSr50 (Chen *et al.,* 2017) and AvrSr35 (Salcedo *et al.,* 2017) are 132 aas and 578 aas long, respectively. With sufficient training data, EffectorP could learn to recognize classes of effectors that share no sequence similarity, yet are structurally conserved, such as MAX-effectors (de Guillen *et al.,* 2015). Machine learning classifiers trained to recognize oomycete RxLR effectors could be used to search for effectors with similar structural properties in fungi. In general, future re-training of EffectorP on the expanding sets of experimentally supported effectors will be critical to retain its value. We envision that in the future separate training sets of apoplastic fungal effectors and cytoplasmic fungal effectors could be of sufficient size to allow for training of separate classifiers, which could potentially increase prediction accuracy. Whilst the machine learning classifier ApoplastP delivers accurate prediction of apoplastic protein localization for both plant and effector proteins (Sperschneider, J. *et al.,* 2017), other signals unique to apoplastic or cytoplasmic effectors might not be fully utilized by EffectorP yet.

Practical recommendations for fungal effector prediction depend on the application. For example, for subsequent experimental validation where time and resources are limited, a stringent effector screening approach might be most appropriate. This could involve taking either EffectorP 2.0 predicted effectors, or effectors predicted by both versions of EffectorP 1.0/2.0 for maximum stringency. For maximum sensitivity, a union of effector candidates predicted by either EffectorP 1.0 or 2.0 can be used, however this will also result in high false positive rates. If *in planta* expression data is available, effectors expressed highly during infection can be prioritized for experimental validation. Another approach would be to select effectors with highest probability, however this has not been tested extensively by us. However, we did observe that during the identification of the *Puccinia graminis f.* sp. *tritici* effector AvrSr50 (Chen *et al.,* 2017), where over 40 candidate genes had to be functionally screened, applying EffectorP 2.0 and ApoplastP (Sperschneider, J. *et al.,* 2017) to predict the most likely effector to enter plant cells would have revealed AvrSr50 as the top candidate with highest probability. Overall, the re-evaluation and re-training of EffectorP has supported the power of machine learning for fungal effector prediction. Higher accuracy of fungal effector prediction will boost experimental validation success rates and aid in the understanding of effector biology.

## Experimental Procedures

### Training of the machine learning classifier

As a positive training set, we collected validated fungal effectors from the literature and then reduced sequence homology in this set by removing those that share similarity with another effector in the set at bit score ≥ 50 using phmmer (Finn *et al.,* 2011). Three negative training sets were generated based on secretomes predicted from annotated gene sets of publicly available genome assemblies of either plant pathogenic fungi and symbionts (21 species, same species from the positive effector training set), animal pathogenic fungi (10 species) or saprophytic fungi (27 species) (Table 7). A protein was labelled as secreted if it is predicted as secreted by the neural network predictor of SignalP 3 (Bendtsen *et al.,* 2004) as well as by TargetP (Emanuelsson *et al.,* 2000) and if it has no predicted transmembrane domain outside the first 60 aas using TMHMM (Krogh *et al.,* 2001), as described previously for fungal effector prediction (Sperschneider *et al.,* 2015b). Each negative set was homology-reduced by deleting proteins that share sequence similarity (bit score ≤ 100, phmmer) with another one in the negative set. We also applied EffectorP 1.0 (Sperschneider *et al.,* 2016) to exclude predicted effectors from the fungal pathogen/symbiont secretomes. The WEKA tool box (version 3.8.1) was used to train and evaluate the performance of different machine learning classifiers (Hall *et al.,* 2009) and feature vectors were calculated for each protein (Table 8). The training data is available at http://effectorp.csiro.au/data.html.

**Table 7:**
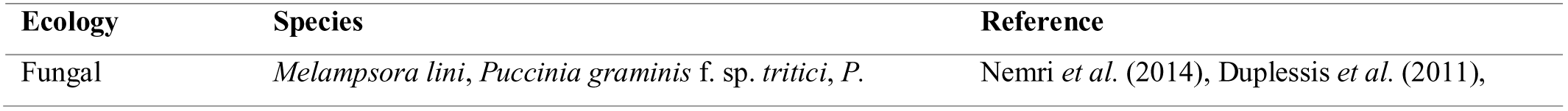
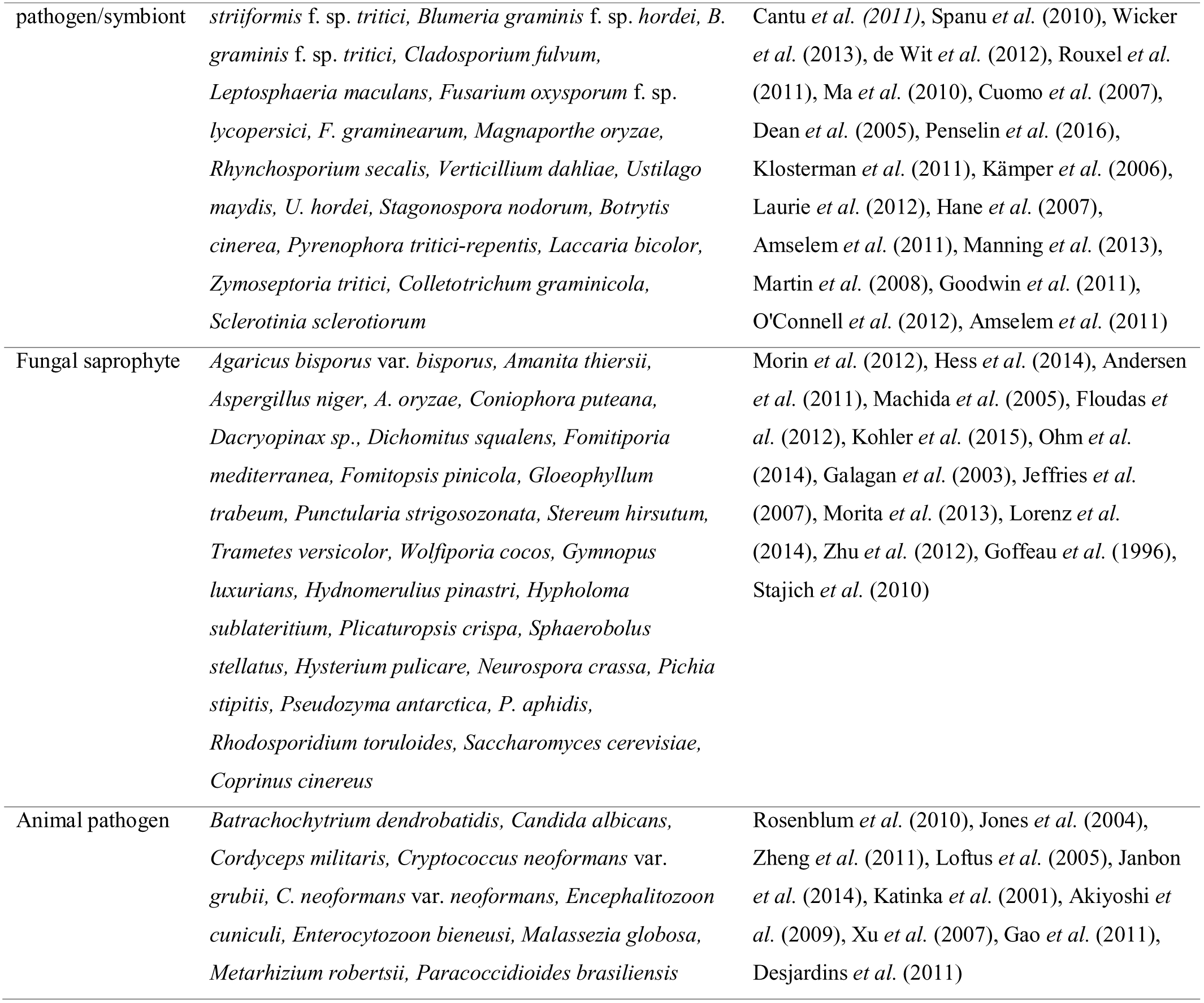
Genomes that were used to predict secretomes for negative training data.

| Ecology | Species | Reference |
| --- | --- | --- |
| Fungal | *Melampsora lini*, *Puccinia graminis* f. sp. *tritici*, *P.* | Nemri *et al.* (2014), Duplessis *et al.* (2011), |
| pathogen/symbiont | *striiformis* f. sp. *tritici*, *Blumeria graminis* f. sp. *hordei*, *B. graminis* f. sp. *tritici*, *Cladosporium fulvum*, *Leptosphaeria maculans*, *Fusarium oxysporum* f. sp. *lycopersici*, *F. graminearum*, *Magnaporthe oryzae*, *Rhynchosporium secalis*, *Verticillium dahliae*, *Ustilago maydis*, *U. hordei*, *Stagonospora nodorum*, *Botrytis cinerea*, *Pyrenophora tritici-repentis*, *Laccaria bicolor*, *Zymoseptoria tritici*, *Colletotrichum graminicola*, *Sclerotinia sclerotiorum* | Cantu *et al.* (2011), Spanu *et al.* (2010), Wicker *et al.* (2013), de Wit *et al.* (2012), Rouxel *et al.* (2011), Ma *et al.* (2010), Cuomo *et al.* (2007), Dean *et al.* (2005), Penselin *et al.* (2016), Klosterman *et al.* (2011), Kämper *et al.* (2006), Laurie *et al.* (2012), Hane *et al.* (2007), Amselem *et al.* (2011), Manning *et al.* (2013), Martin *et al.* (2008), Goodwin *et al.* (2011), O'Connell *et al.* (2012), Amselem *et al.* (2011) |
| Fungal saprophyte | *Agaricus bisporus* var. *bisporus*, *Amanita thiersii*, *Aspergillus niger*, *A. oryzae*, *Coniophora puteana*, *Dacryopinax* sp., *Dichomitus squalens*, *Fomitiporia mediterranea*, *Fomitopsis pinicola*, *Gloeophyllum trabeum*, *Punctularia strigosozonata*, *Stereum hirsutum*, *Trametes versicolor*, *Wolfiporia cocos*, *Gymnopus luxurians*, *Hydnomerulius pinastri*, *Hypholoma sublateritium*, *Plicaturopsis crispa*, *Sphaerobolus stellatus*, *Hysterium pulicare*, *Neurospora crassa*, *Pichia stipitis*, *Pseudozyma antarctica*, *P. aphidis*, *Rhodospiridium toruloides*, *Saccharomyces cerevisiae*, *Coprinus cinereus* | Morin *et al.* (2012), Hess *et al.* (2014), Andersen *et al.* (2011), Machida *et al.* (2005), Floudas *et al.* (2012), Kohler *et al.* (2015), Ohm *et al.* (2014), Galagan *et al.* (2003), Jeffries *et al.* (2007), Morita *et al.* (2013), Lorenz *et al.* (2014), Zhu *et al.* (2012), Goffeau *et al.* (1996), Stajich *et al.* (2010) |
| Animal pathogen | *Batrachochytrium dendrobatidis*, *Candida albicans*, *Cordyceps militaris*, *Cryptococcus neoformans* var. *grubii*, *C. neoformans* var. *neoformans*, *Encephalitozoon cuniculi*, *Enterocytozoon bieneusi*, *Malassezia globosa*, *Metarhizium robertsii*, *Paracoccidioides brasiliensis* | Rosenblum *et al.* (2010), Jones *et al.* (2004), Zheng *et al.* (2011), Loftus *et al.* (2005), Janbon *et al.* (2014), Katinka *et al.* (2001), Akiyoshi *et al.* (2009), Xu *et al.* (2007), Gao *et al.* (2011), Desjardins *et al.* (2011) |

**Table 8:**
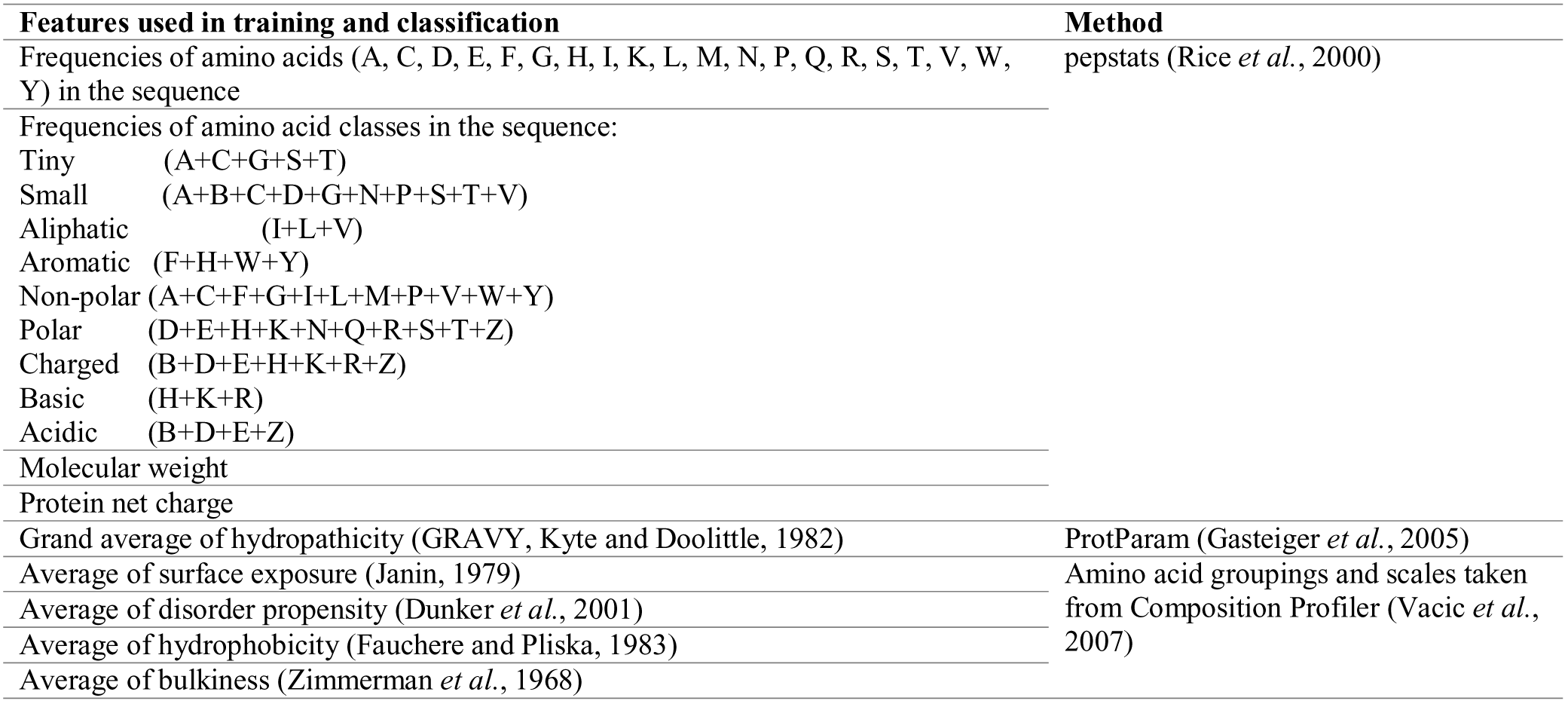
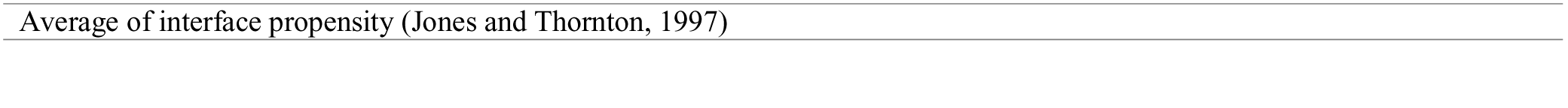
Features used for training the machine learning classifiers in the EffectorP 2.0 ensemble learner.

For the ensemble learner, we took 100 randomly selected samples of negative training data from each of the three negative sets (pathogen/symbiont secretomes, saprophyte secretomes, animal pathogen secretomes), each with 282 protein sequences to give a ratio of 3:1 to the number of positive training examples. We then used WEKA to train Naïve Bayes classifiers on each of the 300 negative data sets with the same positive training set. We then repeated this procedure and trained C4.5 decision trees (J48 model in WEKA) on another 300 randomly chosen negative data from the three classes. For each set of 100 models, we selected the best-performing models as those with the highest area under the curve (AUC). Overall, we chose a total of 50 models comprised of: ten Naïve Bayes classifiers and ten C4.5 decision trees that discriminate between fungal effectors and secreted pathogen proteins; ten Naïve Bayes classifiers and ten C4.5 decision trees that discriminate between fungal effectors and secreted saprophyte proteins; and five Naïve Bayes classifiers and five C4.5 decision trees that discriminate between fungal effectors and secreted animal pathogen proteins. The ensemble classifier called EffectorP 2.0 returns a final prediction using a soft voting approach, which predicts the class label based on average probabilities for ‘effector’ and ‘non-effector’ calculated by each classifier. Soft voting then returns the class with the highest average probability as the result. A protein is classified as an effector if it has probability > 0.55. If it is predicted as an effector with probability 0.5 to 0.55, it is labelled as an ‘unlikely effector’ and is counted as a non-effector in the evaluation.

### Evaluation of EffectorP 2.0

We collected fungal, plant and mammalian proteins with experimentally validated localization to ER, Golgi or membranes or with GPI-anchors from the UniProt database (search terms in Supporting Information Table S1) and predicted signal peptides using SignalP 4.1 (Petersen *et al.,* 2011). We also collected fungal proteins from PHI-base (Urban *et al.,* 2017) from *Fusarium, Magnaporthe, Ustilago, Sclerotinia, Botrytis, Zymoseptoria* and *Leptosphaeria* pathogens that are annotated as having an unaffected pathogenicity phenotype. All evaluation data is available at http://effectorp.csiro.au/data.html.

In the evaluation, true positives (TPs), false positives (FPs), true negatives (TNs) and false negatives (FNs) are calculated. Accuracy is reported as (TP + TN)/(TP + TN + FP + FN), whereas sensitivity is the fraction of effectors that are correctly identified as such [TP/(TP + FN)] and specificity is the fraction of non-effectors which are correctly identified as such [TN/(TN + FP)]. The positive predictive value (PPV) is the proportion of positive results that are true positives [TP/(TP + FP)]. Receiver operating characteristic (ROC) curves plot sensitivity against (1 - specificity) and the area under the curve (AUC) can be calculated. This value gives the probability that a classifier will rank a randomly chosen effector higher than a randomly chosen non-effector. Therefore, a perfect classifier achieves an AUC of 1.0, whereas a random classifier achieves an AUC of only 0.5.

A small size classifier predicts a protein as an effector if it has sequence length of <= 300 aas and a small, cysteine-rich classifier predicts a protein as an effector if it has sequence length of <= 300 aas and >= 4 cysteines in its sequence.

### Functional enrichment analysis and plotting

We performed sequence similarity searches against fungal proteins in NCBI with Blast2GO 4.1.9 (Gotz *et al.,* 2008) and default parameters. GO terms were reduced to the most specific terms and Fisher’s exact tests were used to find over‐ and under-represented terms. Enrichment was called at false discovery rate FDR < 0.05.

Plots were produced using ggplot2 (Wickham, 2009) and statistical significance was assessed with /-tests using the ggsignif package (https://cran.r-project.org/web/packages/ggsignif/index.html). Significance thresholds according to *t-*test are NS: not significant, * < 0.05, ** < 0.01 and *** < 0.001.

## Acknowledgements

J.S. is supported by a CSIRO OCE Postdoctoral Fellowship. We thank Jonathan Anderson and Jonathan Powell for their comments on the manuscript.

## Author Contributions

J.S. planned and designed the research and developed the software. All authors analysed data and wrote the manuscript.

